# Rectal Swab DNA Collection Protocol for PCR Genotyping in Rats

**DOI:** 10.1101/2024.03.02.583131

**Authors:** Audrey E. Kaye, Jacob W. Proctor-Bonbright, Jai Y. Yu

## Abstract

DNA collection is essential for genotyping laboratory animals. However, common collection methods require tissue amputation, causing discomfort and injury. Rectal swabbing has been proposed as an effective non-invasive alternative, but an evidence-backed protocol for the technique remains unavailable. We evaluate the effect of collection parameters on PCR result quality and present a genotyping protocol that can yield results for a litter of rats within 3-5 hours. We found that samples with 2-8 scrapes produced enough DNA to amplify targets up to ∼1800bp long using PCR. Rectal swabbing produced PCR results with similar quality to ear clip samples, and results were unaffected by residual fecal matter or cell debris. Our protocol enables fast, non-invasive, and repeatable genotyping using commercial PCR reagents.

## Introduction

Collecting DNA is essential for genotyping transgenic animal models. Many collection techniques provide DNA samples that can be amplified and analyzed with polymerase chain reaction (PCR). These techniques vary in invasiveness, repeatability, and ease of processing. The most commonly used techniques include ear, tail, and distal phalanx clipping [1], which yield high quantities of DNA but are invasive. Clipping involves the removal or partial amputation of body appendages, causing injury and often requiring a recovery period as well as local or general anesthetics to reduce pain [2-4]. Tail and distal phalanx clipping both have age-related limitations as well, relying on small time windows for sample collection and genotyping [4-7]. Because of their invasiveness and risk for injury, repeated collection with these techniques is either discouraged or not permitted [2,5,8,9]. Alternatively, non-invasive techniques such as fecal pellet collection, oral swabbing, and hair follicle collection [5] can be effective and non-injurious. However, they too pose unique challenges. Fecal pellet collection can be more time-consuming than other non-invasive methods [5,10,11], oral swabbing has a lower expected DNA yield [5], and hair follicle samples can cling to equipment, increasing the risk of cross-contamination [12].

Rectal swabbing [13] is a promising alternative non-invasive collection technique. This method was reported to be quick, easily repeatable, and effective for genotyping both weanlings and adult mice [13]. However, we were unable to find details of the technique in the original report or in the few subsequent published works [5,12]. Thus, there is a lack of information on how to apply the rectal swab technique for DNA collection, and understanding of the critical parameters that determine its success. In this report, we evaluated how collection parameters, such as scrape number, sample number, and fecal matter or cell debris contamination, affected the genotyping outcome for two genomic targets of different lengths: parvalbumin cre (*Pvalb*^*Cre*^, 1803bp) [14], and *Sox21* (237bp). Informed by our findings, we devised a new rectal swab genotyping protocol that repurposes a commercially available PCR kit for quick collection, DNA extraction and PCR. With this protocol, sample collection and PCR genotyping for an average litter of rats can be completed within three to five hours. We observe that rectal swabbing is an effective, non-invasive, and repeatable DNA collection technique for PCR genotyping.

## Materials and Methods

### Experimental Rats and Materials

We evaluated the rectal swab genotyping technique using 12 heterozygous transgenic *Pvalb*^*Cre*^ rats (6, 17, and 27 weeks old, 7 males and 5 females) [14]. DNA extraction and PCR was performed using the Thermo Scientific**™** Phire Tissue Direct PCR Master Mix (#F-170S).

### Rectal Swabbing

Rats were gently restrained and a sterile 1 μL inoculation loop (Fisher Scientific 22-363-601) was inserted into the rat’s rectum. The loop was used to carefully scrape the rectal lining and collect epithelial cells. We waited for any droppings to exit the rectum before inserting the inoculation loop and repeated collection if there was visible fecal matter on the loop. The tip of the loop was then briefly swirled and clipped off into a 0.2 mL tube prefilled with 20 μL of Dilution Buffer from the Phire Tissue Direct PCR kit.

Following manufacturer instructions [15], we added 0.5 μL of DNARelease Additive to each tube before briefly vortexing and centrifuging. The sample were incubated at room temperature (∼20°C) for 2-5 minutes, heated to 98°C for 2 minutes, and then centrifuged for ∼15 seconds as to separate cellular debris from supernatant. 13 μL of supernatant was withdrawn and stored (–20°C), 1μl of which was used for PCR.

### Experimental Variations

We varied the proposed parameters of scrape number, fecal matter, and cell debris to test how they each affect the outcome of the genotyping procedure. To test the effect of scrape number, we collected samples with various scrape numbers (2, 4, 8, 12, 16, and 20) from each rat. To maintain consistency, we used 16 scrapes per sample for subsequent experiments. To test the technique’s reproducibility, we collected three additional samples from each rat. To test the effect of fecal matter, we collected one sample with visible fecal matter on the inoculation loop (n=6, 4 male and 2 female). To test the effect of cellular debris, we collected one sample from each rat but did not centrifuge supernatant away from debris during DNA extraction.

### Ear clipping

We clipped a piece of tissue from 6 experimental rats’ ears (3 males, 3 females) using a 0.5 mm puncher. Then, we extracted DNA using the same procedure and reagents as for the rectal swab samples.

### DNA Quantification

We quantified DNA concentration using 2μL of extracted sample on a Thermo Scientific NanoDrop 1000 Spectrophotometer (ND-1000).

### PCR

The *Pvalb*^*Cre*^ product (1803bp) was amplified using forward primer sequence 5’-GTCATGAACTATATCCGTAACCTGG-3’ and reverse primer sequence 5’-AGTGGTGCACACCCTG ATAC-3’ with final concentrations of 0.5μM. The *Sox21* product (237bp) was amplified using the forward and reverse primers 5’-AGCCCTTGGGGASTTGAATTGCTG-3’ and 5’-GCACTCCAGAGGACAGCRGTGTCAATA-3’ respectively, at a final concentration of 0.125μM. We used the following cycling protocol on a Bio-Rad T100 Thermal Cycler: 98 °C for 5 minutes, followed by 35 cycles of 98 °C for 5 seconds, 64 °C for 5 seconds, and 72 °C for 40 seconds. The final extension was 72 °C for 1 minute. Gel electrophoresis was performed using 1% TBE agarose gels with ethidium bromide (25 minutes at 100V, Horizontal Mini Gel Electrophoresis System, Fisher Scientific). PCR results were imaged with a Syngene GeneFlash Gel Imaging System.

### Image quantification

Relative band brightness was quantified using ImageJ. For each band, we defined an 18x20 pixel region of interest (ROI) around the band itself and another around the lane background immediately below it. We calculated relative band brightness as a ratio of the band ROI mean intensity over the background ROI mean intensity.

### Data analysis

We analyzed the data using Python and Jupyter Notebook. Statistical analyses were performed using the Scipy package (https://scipy.org/).

## Results

### Scrape Number

Rectal swabbing requires gently scraping the rectum lining, but the number of scrapes needed per sample was not specified in the original report [13]. Institutional guides recommend different ranges of scrapes be made [16], but it is unknown if or how these values have been evaluated for efficacy. We tested how the number of scrapes affects the resulting sample’s DNA concentration and the quality of its PCR amplification outcome. Given that longer PCR products can be challenging to amplify with crude DNA preparations, we evaluated the effectiveness of the procedure on two products of differing length: *Sox21* (273bp), and *Pvalb*^*Cre*^ (1803bp).

We anticipated that more scrapes would create higher concentration samples and brighter bands on PCR, but found otherwise. Instead, samples from 2 to 20 scrapes all yielded enough DNA to produce visible and comparably bright bands on a first-round PCR. The scrape number used for collection was not correlated with DNA concentration (Fig 2B, r^2^ = 0.0009 and p = 0.80). Similarly, scrape number was not correlated with the brightness of the *Pvalb*^*Cre*^ PCR product (Fig. 2C, r^2^ = 0.0067 and p = 0.50) or the *Sox21* product (Fig. 2D, r^2^ = 0.0019 and p = 0.72). This suggests that any number of scrapes from 2 to 20 can collect sufficient DNA for PCR amplification. We therefore recommend that at least 2-8 scrapes be completed per sample.

**Figure 1.**
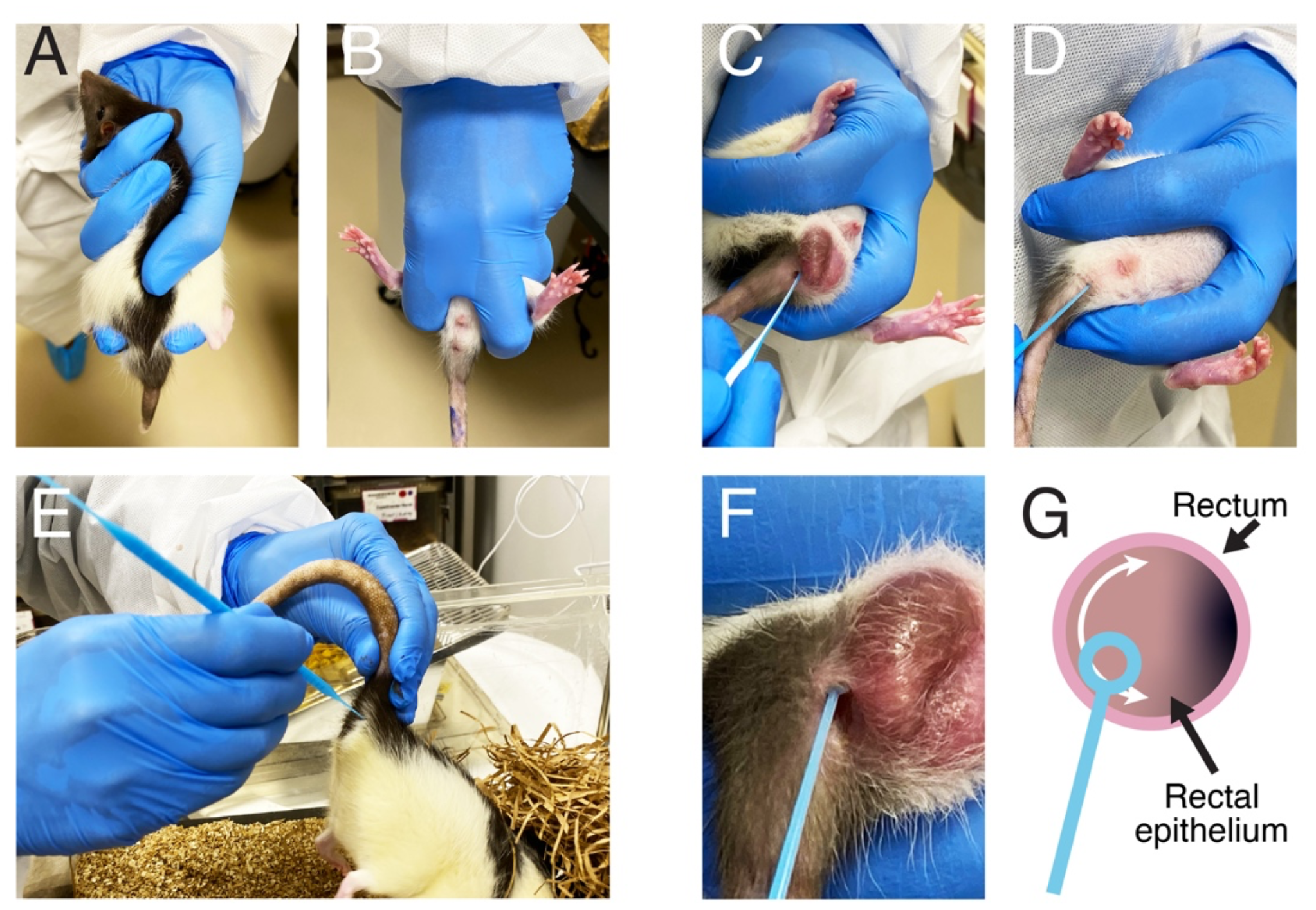
Hold technique for rectal swab collection. **(A)** Dorsal view demonstrating hold for young or small rats (<200g). **(B)** Ventral view of hold for young or small rats. **(C)** Positioning the inoculating loop for collection in small male rats. **(D)** Positioning the inoculating loop for collection in small female rats. **(E)** Hold position for large rats (>200g). **(F)** Close-up view of loop insertion. **(G)** Schematic of scraping technique.

**Figure 2.**
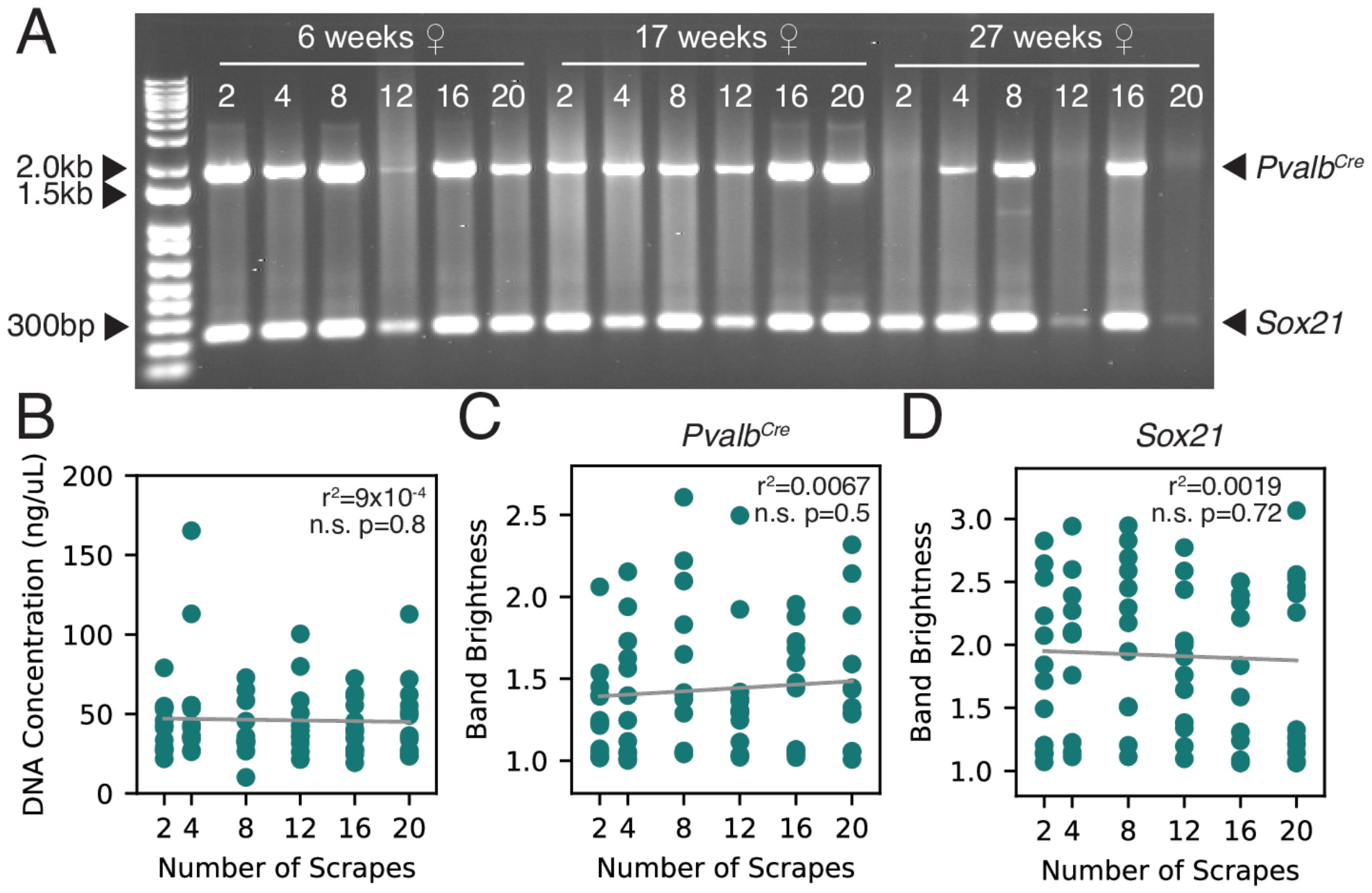
PCR outcome and DNA concentration does not depend on the number of scrapes used for collection. **(A)** Example PCR gel image with samples of varying scrape number collected from three rats of varying age. **(B)** Scatter plot of sample DNA concentration (ng/uL) versus scrape number (n.s. r^2^ = 0.00090 and p = 0.80). **(C)** Scatter plot of *Pvalb*^*Cre*^ PCR product band brightness versus rectal scrape number (n.s. r^2^ = 0.0067 and p = 0.50). **(D)** Scatter plot of *Sox21* PCR product band brightness versus rectal scrape number (n.s. r^2^ = 0.0019 and p = 0.72). Statistical testing was conducted using a linear regression test.

### Sample Number

To determine the optimal number of samples per rat necessary for producing reliable results, we first performed PCR on four rectal swab samples from each known-positive animal (n=12) and evaluated whether the *Pvalb*^*Cre*^ and the *Sox21* products were amplified consistently. We observed both *Pvalb*^*Cre*^ (1803bp) and *Sox21* (273bp) product bands for most PCRs.

However, some reactions only produced the *Sox21* band and not the *Pvalb*^*Cre*^ band. Given that the *Pvalb*^*Cre*^ product is longer than the *Sox21* product, it may be more difficult to amplify due to potential degradation of longer strands in the crude DNA mixture. Therefore, while the rectal swab technique reliably collected sufficient DNA for amplification, different length target sequences may vary in amplification success during PCR.

Due to this variability, we asked how many samples are needed from each animal to maximize the likelihood of detecting a true positive PCR result. With statistical sampling, we simulated the effect of taking 1-4 samples from each of the 12 positive rats on the likelihood of observing a positive PCR outcome. For instance, we simulated a collection with two samples per rat by randomly selecting two of their four experimentally observed outcomes. After doing this for all 12 rats, we defined the detection rate as the proportion of rats that had at least one positive result. We then repeated this simulation 10,000 times and calculated the average detection rate.

Our simulations indicate that one rectal swab sample is sufficient to produce a visible PCR band 75% and 94% of the time for *Pvalb*^*Cre*^ and *Sox21* respectively (Fig. 3). However, taking two samples per animal would increase this success rate to 89% for *Pvalb*^*Cre*^ and 97% for *Sox21*. We saw the largest increase in amplification success between one and two samples (14.52% for *Pvalb*^*Cre*^ and 3.55% for *Sox21*) followed by marginal improvement with every additional sample taken (2^nd^-3^rd^: 5.98% and 3^rd^-4^th^: 2.34% for *Pvalb*^*Cre*^, 2^nd^-3^rd^: 1.48% and 3^rd^-4^th^: 0.63% for *Sox21*). As such, we recommend taking two samples per rat to maximize the probability of detecting a true positive PCR result with the least additional effort.

**Figure 3.**
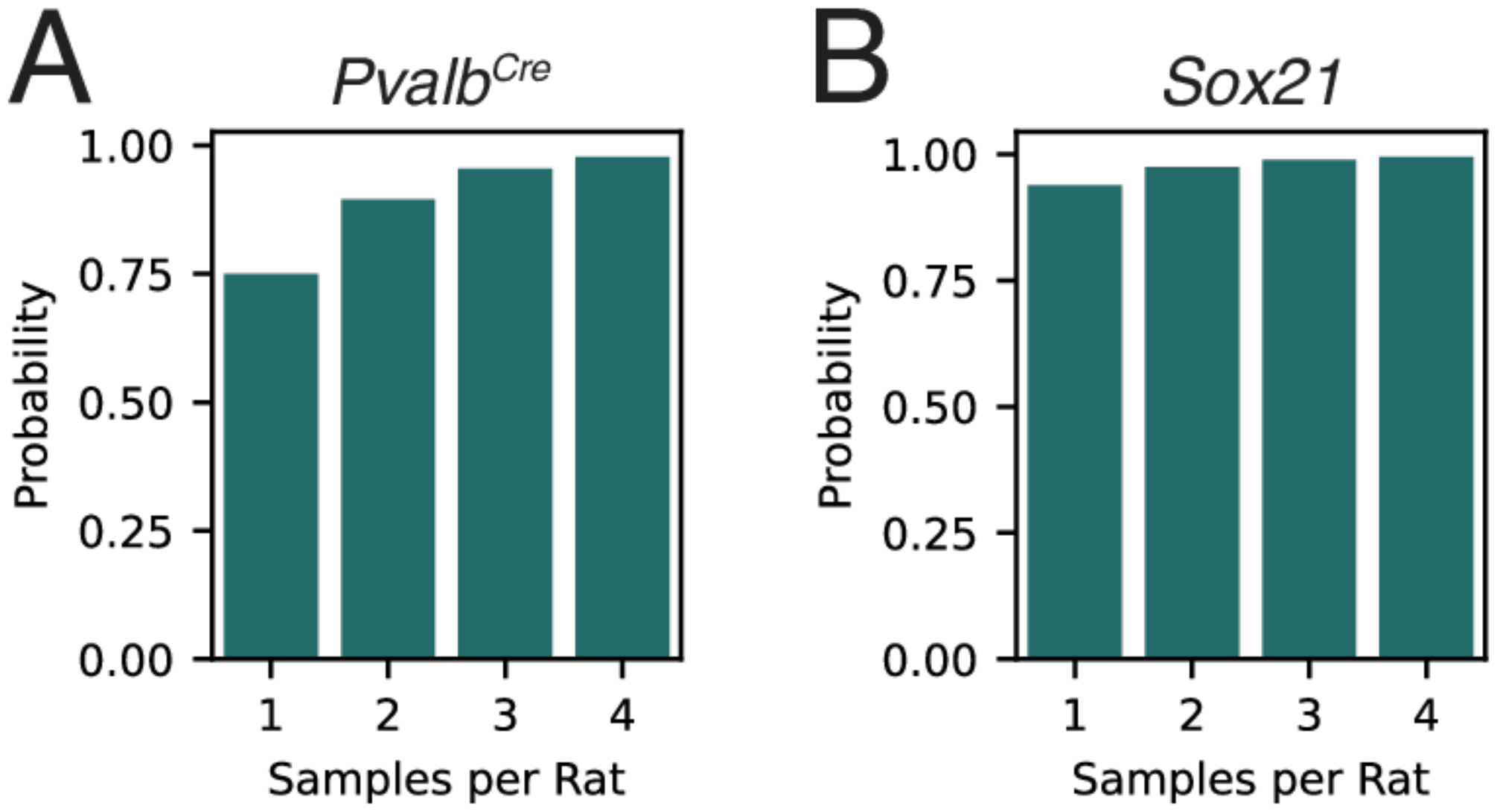
**(A)** Bar plot of the probability of observing least one visible *Pvalb*^*Cre*^ PCR product band given the number of samples collected from a known-positive *Pvalb*^*Cre*^ rat. **(B)** Bar plot of the probability of observing least one visible *Sox21* PCR product band given the number of samples collected.

### Fecal Matter & Cellular Debris

Two potential sources of contamination during this procedure are fecal matter from sample collection and cell debris from DNA extraction. Fecal matter contains gut microflora DNA and other compounds that interfere with the PCR process, such as bile salts, complex polysaccharides, and hemoglobin products [10,17,18]. Additionally, PCR guidelines specify that cell debris be separated from DNA supernatant to prevent unwanted particles from interfering with results. For these reasons, we tested whether collection should be repeated in cases where fecal matter or cell debris remain in the sample.

To do this, we tested how fecal matter and cell debris affect PCR success. We found that, while the presence of fecal matter significantly increased sample DNA concentration (Fig. 4B, p = 0.031), it did not significantly affect brightness of the resulting *Sox21* band (Fig. 4D, p = 1.0) or *Pvalb*^*Cre*^ band (Fig. 4C, p = 0.31) on PCR. Because fecal matter increases DNA concentration without increasing band brightness, the additional DNA likely comes from rectal microflora. As fecal matter did not affect PCR results, small amounts of feces on the collection loop does not warrant repeating the collection. Additionally, uncentrifuged cell debris did not significantly affect DNA concentration (Fig. 4E, p = 0.69), *Sox21* band brightness (Fig. 4G, p = 0.84), or *Pvalb*^*Cre*^ band brightness (Fig. 4F, p = 0.22) of the resulting sample. Thus, incompletely centrifuged cell debris does not require sample recollection.

**Figure 4.**
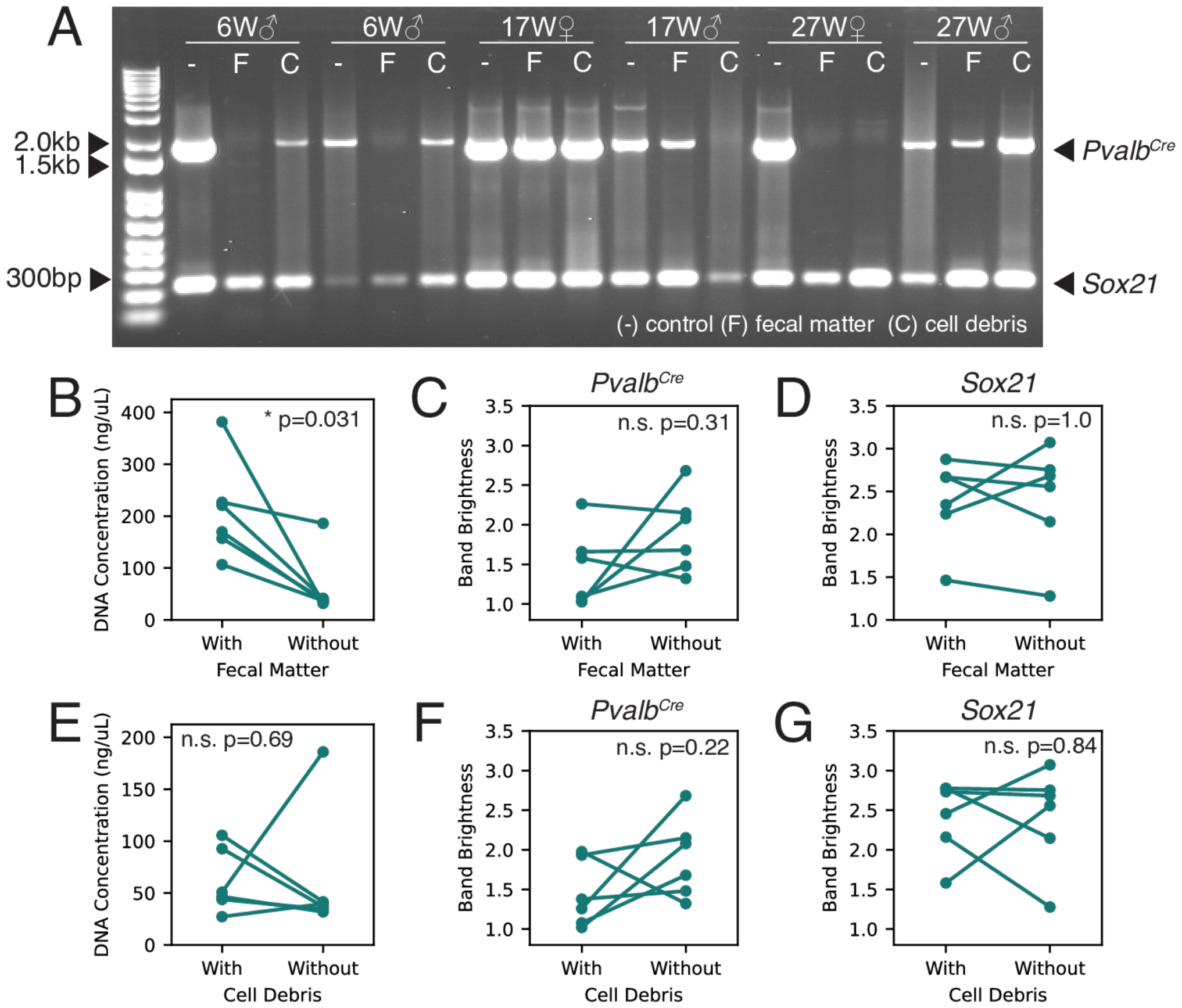
Fecal matter increases DNA concentration but does not affect PCR amplification, while cell debris presence affects neither metric. **(A)** Sample PCR including proper rectal swab samples, samples with visible fecal matter, and samples with persistent cell debris from rats C, D, E, H, J, and L. **(B)** Scatter plot of DNA concentration versus fecal matter presence for six rats of varying ages (p = 0.031). **(C)** Scatter plot of *Pvalb*^*Cre*^ band brightness on PCR versus fecal matter presence (n.s. p = 0.31). **(D)** Scatter plot of *Sox21* band brightness on PCR versus fecal matter presence (n.s. p = 1.0). **(E)** Scatter plot of sample DNA concentration versus cell debris presence (n.s. p = 0.69). **(F)** Scatter plot of *Pvalb*^*Cre*^ band brightness on PCR versus cell debris presence (n.s. p = 0.22). **(G)** Scatter plot of *Sox21* band brightness on PCR versus cell debris presence (n.s. p = 0.84). Statistical testing was conducted using the Wilcoxon signed-rank test.

**Figure 5.**
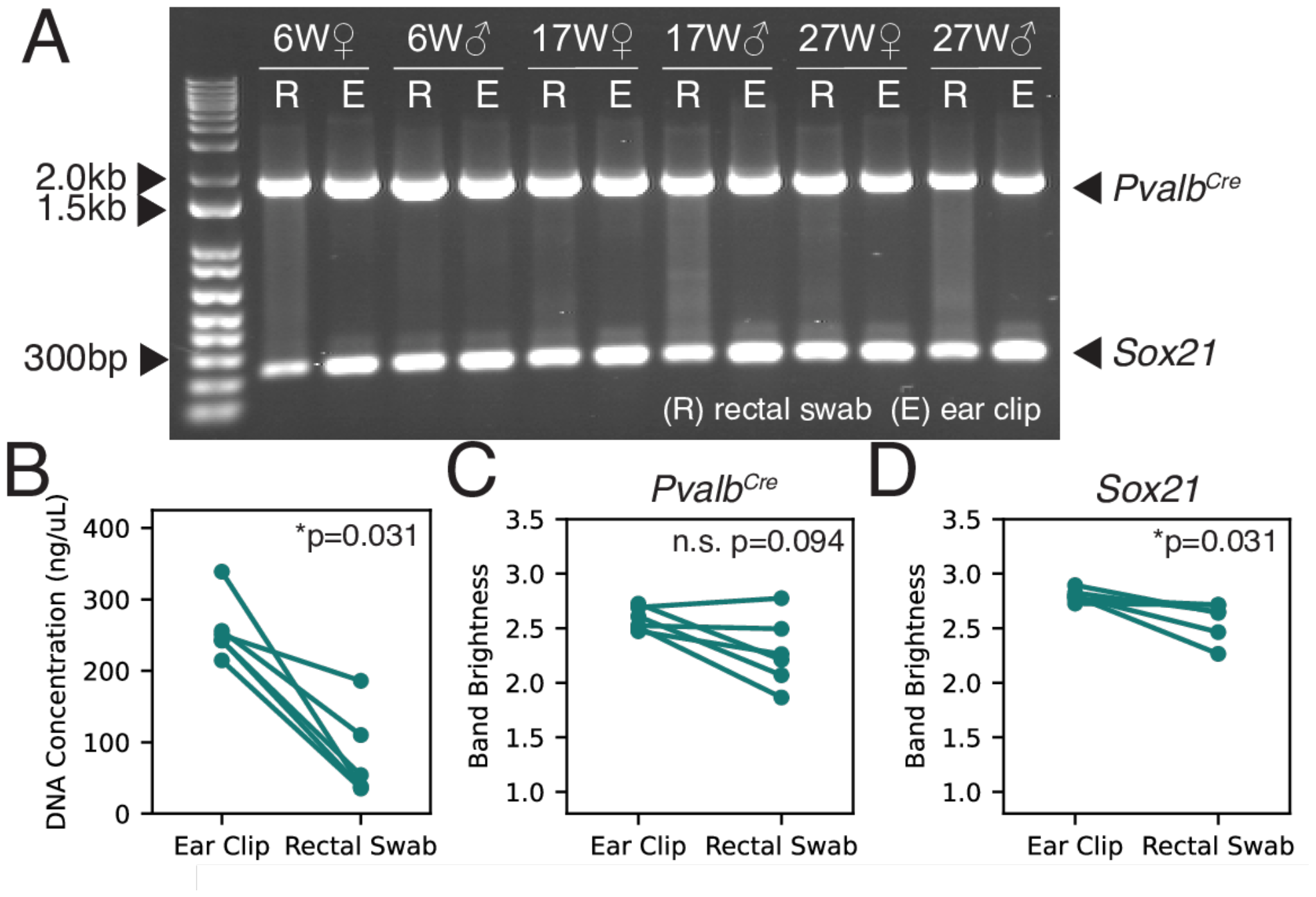
Ear clipping produced higher DNA concentration and brighter *Sox21* bands than rectal swabbing, but both techniques produce *Pvalb*^*Cre*^ bands of statistically comparable brightness. **(A)** Sample gel including PCR-amplified rectal swab and ear clip samples from six rats of varying ages and sexes. **(B)** Paired scatter plot of DNA concentration versus DNA collection technique (p = 0.031). **(C)** Paired scatter plot of *Pvalb*^*Cre*^ band brightness on PCR versus DNA collection technique (n.s. p = 0.094). **(D)** Paired scatter plot of *Sox21* band brightness on PCR versus DNA collection technique (p = 0.031). Statistical testing was conducted using the Wilcoxon signed-rank test.

### Ear Clip Genotyping

We compared rectal swabbing to the common yet invasive ear clipping technique. We found that ear clipping produced significantly higher DNA concentrations (Fig. 4B, p = 0.031) and brighter *Sox21* bands (Fig. 4D, p = 0.031), but produced no significant difference in *Pvalb*^*Cre*^ band brightness (Fig. 4C, p = 0.094). Despite differences in band brightness, both techniques produced visible bands that clearly convey the genotype of a transgenic animal. Thus, rectal swabbing can produce DNA of sufficient quality for use in PCR reactions, comparable to the established method of ear clipping.

## Discussion and Conclusion

Our results indicate that the rectal swab technique is effective for collecting sufficient genomic DNA for PCR amplification. The number of scrapes made along the rectal epithelium did not affect PCR outcome quality, allowing us to recommend a lower range of 2-8 scrapes per sample. Taking two samples per rodent further increased the likelihood of PCR success.

Additionally, fecal matter and cell debris did not affect PCR outcome either, indicating that collection need not be repeated in these cases. While ear clipping produced higher DNA concentrations, rectal swabbing nonetheless collected sufficient DNA to produce visible PCR bands for sequences up to ∼1800bp. Finally, our protocol leverages the efficiency of commercial direct-from-tissue PCR reagents to enable DNA extraction and PCR with a time frame and cost that are comparable to invasively collected tissue samples. We conclude that rectal swabbing is similarly effective for genotyping rodents with PCR, while remaining non-invasive, repeatable, and comparably quick and inexpensive.

The complete process of rectal swab genotyping, including sample collection, PCR, and gel electrophoresis, takes approximately 3-5 hours for a typical litter of laboratory rats with 8-16 pups. We note that the previous protocol outlined by Lahm, et al. used overnight DNA extraction [13], while our extraction process takes approximately 5-10 minutes for one sample. One report claims that rectal swab genotyping requires more reagents [19], however, our protocol uses the a set of reagents and manufacturer guidelines designed to process tissue samples. Our use of these commercial kits demonstrates that rectal swab samples can be processed at the same speed and cost per reaction as invasively-collected tissue samples.

Finally, the rectal swab technique is also advantageous for animal welfare. It is a non-invasive technique that does not require amputation like more popular invasive methods. Physiological measurements indicate that rectal swabs do not cause long-term distress to animals [12]. For these reasons, rectal swabbing can be repeated many times without injuring the rat or running out of tissue for collection, which is possible for many invasive techniques. Tail clipping, for instance, can only be completed twice, with the second sample being highly discouraged [8,9]. One reported concern is bleeding during rectal swabbing in mice [12,16], which implies injury and thus raises the question of invasiveness. However, we have not observed any bleeding with adult or weaning-aged rats while using this technique. We report that rectal swabbing can be conducted safely and effectively without causing injury.

### Future Perspective

Non-invasive DNA collection is important for minimizing distress to animals from injury or amputation. This is especially important for studies investigating locomotion and fine limb movement. Additionally, fast and convenient genotyping methods will be highly suitable for laboratories with minimal molecular biology equipment. Our protocol can potentially be applied to laboratory species other than rats and mice, including gerbils, hamsters, and guinea pigs. PCR genotyping is used in these model species to determine natural genetic variation or for sexing.

Further, rectal swabbing may be used to collect DNA from small animals in field studies where fecal samples are difficult to collect, or where tissue sampling through blood or appendage amputation poses even higher risk of harm. Our protocol thus provides researchers with a convenient method for DNA collection with a wide range of applications in and out of the laboratory.

## Procedure

We devised the following protocol for rectal swab collection, processing, and PCR analysis based on our evaluation of scrape number, reproducibility, and possible contaminants.

### 1. Secure the rat gently but firmly

For smaller rats (<200g), place the rat’s belly on your palm tail-side outward and hook the rear with your index and middle finger on either side of the tail. From the dorsal side, secure the torso with your thumb as well as the head and neck with your ring and pinky fingers (Fig. 1A-B). Hold the rat ventral side up for visibility during collection. For rats larger than the size of your hand, hold the rat by the base of the tail and lift its hind legs from behind (Fig. 1E).

### 2. Scrape the rectal lining with a sterile inoculating loop

After the rat is secure, insert a 1uL inoculation loop into the rectum, in line with the direction of the rat’s torso (Fig. 1C-D). With the loop flat against the side of the rectal lining, gently scrape it along the epithelium in a semicircular motion 2-8 times, and withdraw (Fig. 1F-G, 2). It is preferable to allow the animal to defecate before inserting the loop into the rectum. However, small amounts of fecal matter on the inoculation loop will not affect the outcome of the procedure and collection need not be redone in that instance (Fig. 4B-D). It is recommended that you collect two samples per animal (Fig. 3). At no point in this process should there be blood on the loop. If you observe blood, you should reduce the force applied.

### 3. Clip the inoculation loop into a collection tube

Submerge the inoculation loop end into a 0.2 mL tube prefilled with 20 uL of Dilution Buffer and briefly swirl. Clip the tip of the loop with metal snips to release it into the tube. Clean the metal snips with alcohol wipes before the next collection, including between samples from the same rat.

### 4. Extract DNA with direct-from-tissue PCR reagents

Extract DNA according to the Thermo Scientific**™** Phire Tissue Direct PCR Master Mix manual [15]. Add 0.5 uL of DNARelease Additive to each tube, briefly vortex and centrifuge the samples, and then let the reaction incubate at room temperature for 2-5 minutes. Heat the samples to 98°C for 2 minutes in a thermocycler, then centrifuge samples for 15-20 seconds to separate cellular debris from supernatant. Remove 10-15 uL of supernatant into a separate 0.2 uL tube for immediate use in PCR or for storage at –20 C. Small amounts of residual debris transferred with the supernatant will not affect the genotyping results (Fig. 4E-G).

### 5. Amplify sample DNA with polymerase chain reactions (PCR)

Complete PCR according to the Thermo Scientific Phire Tissue Direct PCR Master Mix manual [15]. Samples may be stored at –20 C between the PCR and gel electrophoresis.

### 6. Complete gel electrophoresis and image analysis

Run extracted and amplified DNA samples through gel electrophoresis, then photograph for image analysis.

